# Effects of Disease on Renal Blood Flow Regulation: Assessing Autoregulation Disruption

**DOI:** 10.1101/2024.12.09.627590

**Authors:** Bhavishya Maddineni, Ashlee N. Ford Versypt

## Abstract

The functioning of kidneys on blood flow regulation was investigated, particularly under diseased conditions such as chronic kidney disease, which includes conditions of diabetic nephropathy and other glomerular damage. A mathematical model was developed to better understand how variations based on the glomerular filtration rate impact key kidney function outputs, such as afferent arteriolar diameter, smooth muscle activation, and the chloride ion concentration at the macula densa. We have analyzed these factors by considering the dynamics of the mathematical model of ordinary and partial differential equations to study blood flow control in the kidney, which has provided new insights into the maintenance of autoregulation. By simulating the processes of renal blood flow—specifically through the afferent arteriole and glomerulus, and detailing the process of chloride transport within the renal tubule—the model offers a comprehensive view of how the kidney regulates glomerular filtration rate amidst fluctuating systemic blood pressures and disease-specific changes. Central to this model are the myogenic response that adjusts afferent arteriole muscle tone in reaction to pressure changes and the tubuloglomerular feedback, which controls arteriole size based on chloride levels at the macula densa. The model’s simulations reveal the robustness of renal autoregulation across a spectrum of chronic kidney disease stages, showing stability under normal conditions but indicating a breakdown in regulation with advanced chronic kidney disease. This breakdown is attributed to disruptions in the vascular and feedback systems. The findings from this model shed light on the progression of renal dysfunction in chronic kidney disease and underscore the potential for developing targeted treatments to maintain kidney health.

## 1 Introduction

The kidneys are essential organs tasked with the critical role of maintaining bodily homeostasis, a function achieved through the filtration of blood and the production of urine. This process involves the removal of metabolic by-products like urea, creatinine, and uric acid, alongside the regulation of electrolyte balance, blood pressure, acid-base equilibrium, and gluconeogenesis [1]. These tasks are executed by the kidney’s nephrons, numbering approximately one million per organ, which filter blood to produce urine, adjusting the body’s fluid and electrolyte levels through reabsorption and secretion.

Nephrons, the kidney’s fundamental structural and functional units, are composed of the glomerulus, where blood filtration occurs; the juxtaglomerular apparatus, regulating filtration rates; and the renal tubules, transforming filtrate into urine [2]. The kidney is anatomically structured into the cortex, outer medulla, and inner medulla, with the medulla crucial in concentrating urine [3]. Afferent arterioles in the renal cortex maintain nephron blood supply, ensuring efficient filtration and overall homeostasis.

A pivotal indicator of kidney health is the glomerular filtration rate (GFR), a measure of how much blood the kidneys filter per minute, with healthy individuals typically exhibiting a GFR of about 125 mL/min or 180 L/day. Remarkably, of this filtered volume, 99% is reabsorbed into the bloodstream, leaving only 1–2 L to be excreted as urine each day [4]. The kidney’s ability to autoregulate blood flow, crucial for maintaining stable renal function despite systemic blood pressure variations, is central to this process [5]. This autoregulation is achieved through mechanisms such as the myogenic response and tubuloglomerular feedback (TGF). The myogenic response [6] enables afferent arterioles to constrict or dilate in response to changes in blood pressure, stabilizing renal blood flow and GFR. TGF, on the other hand, adjusts GFR based on sodium chloride concentrations sensed by the macula densa in the distal tubule, ensuring homeostasis of fluid and electrolyte excretion [7]. Together, these mechanisms protect the kidneys from pressure-induced damage and maintain optimal functionality.

Chronic kidney disease (CKD) disrupts these autoregulatory mechanisms, with progressive damage impairing the kidney’s ability to maintain hemodynamic stability. CKD is characterized by structural and functional damage to the nephron, manifesting as glomerular sclerosis, tubular atrophy, and interstitial fibrosis, which collectively impair GFR. In advanced stages, CKD severely compromises autoregulatory capacity, predisposing the kidney to further injury and accelerating disease progression. Hypertension, diabetes, and other systemic conditions exacerbate these effects by imposing additional stress on the autoregulatory processes, leading to further deterioration in kidney function.

This study builds upon existing mathematical models of renal autoregulation, particularly those developed by Arciero et al. [8] and Ford Versypt et al. [9], to analyze the impact of CKD on renal hemodynamics and autoregulatory efficacy. By integrating clinical levels of changes into single nephron GFR in these models, we aim to elucidate the changes in autoregulatory function associated with CKD progression. Understanding these dynamics is crucial for developing therapeutic strategies aimed at preserving kidney function and delaying disease progression.

## 2 Methods

Adapted a previously published mathematical model to simulate glomerular filtration rate (GFR) changes and predict their effects on key renal variables, such as vessel diameter and chloride ion (Cl^−^) concentration. Building upon the existing mathematical models by our lab in earlier studies [8] and [9], this model highlights the significant role of the afferent arteriole in regulating renal blood flow. In this model, the afferent arteriole is represented as a single resistor, following Ohm’s law, with its dynamics governed by a system of ordinary differential equations (ODEs). The afferent arteriole’s behavior, influenced by the vessel wall’s mechanical properties, serves as the primary regulator of blood flow into the nephron [8, 9].

Key nephron components, including the glomerulus, proximal tubule, loop of Henle, and descending limb, were integrated into the model, alongside a partial differential equation (PDE) to describe Cl^−^ transport along the thick ascending limb (TAL). The model also incorporates autoregulatory mechanisms, such as the myogenic response and tubuloglomerular feedback (TGF), to simulate the kidney’s regulation of blood flow.

To extend the model for chronic kidney disease (CKD) stages, we calculated single nephron GFR (SNGFR) values based on clinical GFR ranges, using newly developed equations. These values were coupled with the existing model equations to simulate the effects of CKD on renal blood flow dynamics, integrating both myogenic and TGF responses for a comprehensive analysis.

### 2.1 Existing mathematical model

The model was developed based on fluid dynamics and mass transport principles, incorporating the physiological structure of the nephron [8, 9] (Fig. 1). The complexities of renal homeostasis were simplified by assuming a baseline steady-state condition for the flow and solute concentration in the absence of pressure disturbances or flow rate set point changes (corresponding to disease states).

**Fig 1.**
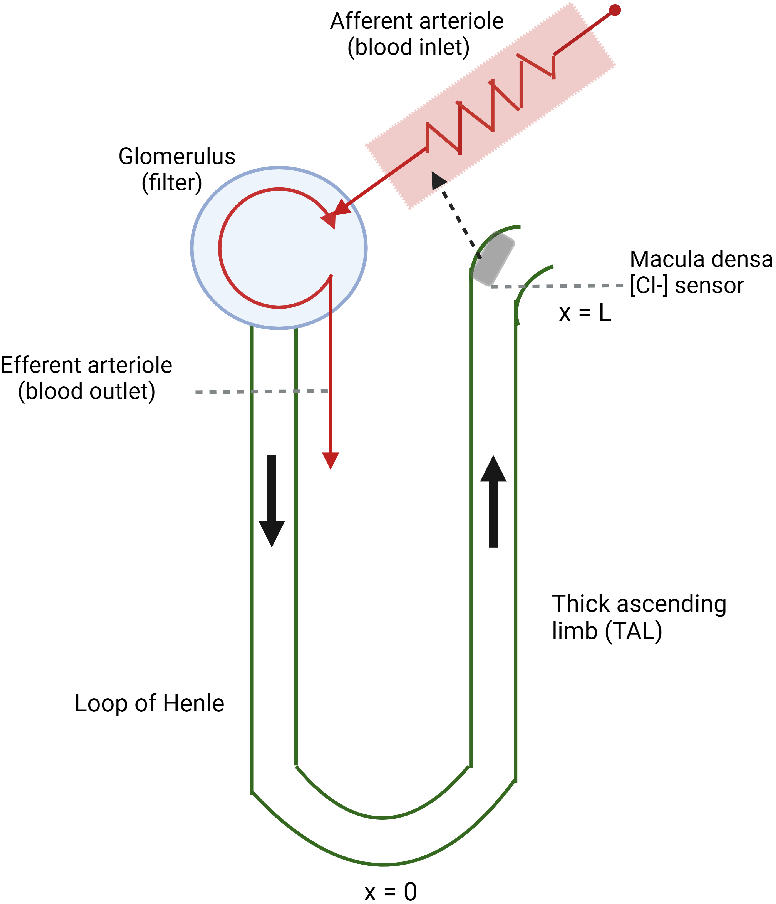
Schematic of single kidney nephron inspired by [8]. Created with BioRender.com.

Here, we provide a brief overview of the existing model. ODEs are used for the arteriole dynamics for the afferent arteriolar diameter and activation smooth muscle tone. These dynamics include the arteriole’s myogenic responsiveness to pressure perturbations, encapsulating the myogenic mechanism and feedback from the TGF mechanism in the nephron’s distal segment. The coupling of these components within the model accounts for the combined effects of both myogenic and TGF responses. The model also features a PDE that captures transport phenomena in the TAL of the loop of Henle. This PDE accounts for diffusion and active transport processes establishing the chloride concentration gradient. The model structure includes the following components:

- Afferent arteriole dynamic responses

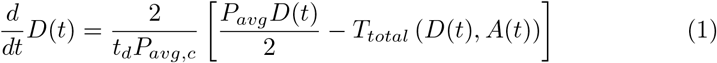

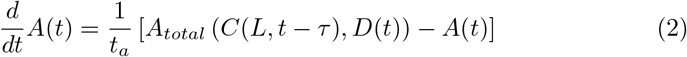

where *t*_*d*_ and *t*_*a*_ are time constants for diameter and activation, respectively, *P*_*avg*_ is the average pressure, the parameter *P*_avg,c_ corresponds to the average pressure when the incoming pressure *P* is at a control state of 100 mm Hg, *T*_*total*_ is the total wall tension that is a nonlinear function of *D* and *A* defined in [8] and [9], *A*_*total*_ is the total possible vascular smooth muscle activation that is a nonlinear function of *A* and *C*(*L, t − τ*) defined in [8] and [9], *A* is the level of activation of the smooth muscles of the blood vessel wall in the afferent arteriole, and *C*(*L, t − τ*) is the Cl^−^ concentration in the TAL at *x* = *L* at the upper end of the TAL at the MD evaluated at a time delayed by *τ* . *C*(*x, t*) is defined in the next model portion in Equation (4). The term *P*_avg_ denotes the average pressure within the afferent arteriole and is calculated based on the incoming pressure and the subsequent pressure drop along the arteriole segment such that

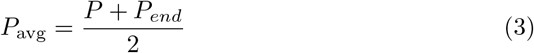

where *P* is the incoming intraluminal pressure and *P*_*end*_ is the pressure at the end of the arteriole segment, which is fixed to 50 mm Hg. For this study, *P* is maintained at 100 mm Hg when not considering pressure disturbances. Alternatively, we also consider variations in *P* from 60 to 180 mm Hg as in [8] to explore impacts of disease conditions on the autoregulatory capacity to reject pressure disturbances.
- Cl^−^ transport in thick ascending limb

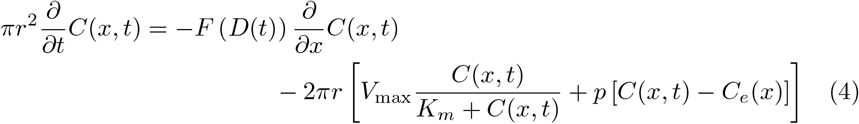

where *C*(*x, t*) represents the Cl^−^ concentration in the TAL at position *x* and time *t*, with *x* = 0 at the bend in the loop of Henle and *x* = *L* at the upper end of the TAL at the MD Fig. 1. *C*_*e*_(*x*) is extratubular Cl^−^ concentration that is defined by a nonlinear equation in [8] and [9]. All other parameters are defined in The equation is derived based on the conservation of mass [8], where the left-hand side in Equation (4) represents the change in Cl^−^ concentration in cylindrical geometry. The right-hand side of Equation (4) accounts for three distinct processes: the axial transport of due to advection, the active transport of solutes out of the tubule, and the diffusion of Cl^−^ across the tubular epithelium.
- Filtrate flow rate The filtrate flow rate, denoted as *F* (*D*(*t*)), is a key component in the advective transport term of Equation (4) and couples the Cl^−^ transport to the afferent arteriole dynamic responses. The filtrate flow rate is explicitly defined as

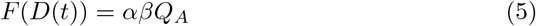

where *α* is the fraction of the SNGFR that is not reabsorbed in the proximal tubule or the descending limb of the loop of Henle before reaching the TAL, *β* represents the fraction of the afferent arteriole flow entering the loop of Henle, and *Q*_*A*_ is the volumetric flow rate defined by the Hagen-Poiseuille equation:

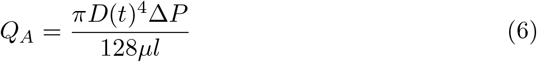

where Δ*P* is the pressure drop along the afferent arteriole (*P − P*_*end*_), *µ* is viscosity of the blood, and *l* is the afferent arteriole segment length. It is important to note that the TAL is assumed to be impermeable to water, which means that the fluid flow along it remains constant in space, though it may vary over time.

The integration of these components forms a comprehensive system of equations. The arteriole dynamic responses govern the filtrate flow rate, which influences the Cl^−^ transport equation. The latter computes the Cl^−^ concentration at the terminal segment of the loop of Henle, which is then coupled back to the arteriole dynamic responses after a time delay, thereby completing the feedback loop. Equations (1)–(4) and parameters are specifically tailored to model the physiological responses to stimuli within the kidney’s regulatory mechanisms. By simulating the interactions between the myogenic response and tubuloglomerular feedback, the model provides insights into how these mechanisms collectively regulate blood flow dynamics in response to changing physiological conditions. The initial and boundary conditions and other detailed derivations are described in [8] and [9]. The parameters are listed in Table 1.

**Table 1.** Values of parameters appearing in the main model. Abbreviations: thick ascending limb (TAL), chloride ion concentration (Cl^−^), macula densa (MD), and afferent arteriole (AA).

| Symbol | Description | Value | References |
| --- | --- | --- | --- |
| $r$ | TAL luminal radius (cm) | $10 \times 10^{-4}$ | [10] |
| $V_{max}$ | Maximum active transport rate ($\text{mol}/\text{cm}^2\text{s}$) | $14.5 \times 10^{-9}$ | [10] |
| $K_m$ | Michaelis constant (mM) | 70.0 | [10] |
| $p$ | TAL $\text{Cl}^-$ permeability (cm/s) | $1.5 \times 10^{-5}$ | [10] |
| $L$ | Length of TAL (cm) | 0.5 | [11] |
| $C_0$ | Luminal $\text{Cl}^-$ at loop of Henle (mM) | 275.0 | [10] |
| $C_e(L)$ | Interstitial $\text{Cl}^-$ at MD (mM) | 150.0 | [10] |
| $t_d$ | Time constant for AA diameter response (s) | 1.0 | [12] |
| $P_{avg,c}$ | Midpoint pressure in AA at control state $P = 100$ mm Hg (mm Hg) | 75 | [13] |
| $t_a$ | Time constant for activation response (s) | 10.0 | [12] |
| $\tau$ | Time delay for MD to AA response (s) | 4.0 | [10] |
| $\beta$ | Glomerular filtration fraction | - | simulated |
| $\alpha$ | Fraction of glomerular filtrate entering TAL | 0.2 | [14] |
| $P_{end}$ | Pressure at end of AA segment (mm Hg) | 50 | [13] |
| $\mu$ | Blood viscosity in AA (cP) | 4.14 | [15] |
| $l$ | AA segment length (cm) | 0.031 | [16] |
| $D_0$ | Passive AA diameter ($\mu\text{m}$) | 33 | calculated |

Table notes: Values and parameters are derived from the specified references, tailored to simulate the physiological conditions of the nephron model.

### 2.2 Incorporating disease-relevant GFR changes into the existing model

The decline in GFR is a critical indicator of kidney function deterioration. Understanding the stages of GFR changes and their impact on renal physiology is essential for early diagnosis and treatment of kidney diseases. This research addresses the gap in knowledge regarding the quantitative relationship between GFR changes and kidney conditions, particularly in CKD. In [8], the model simulations involve modulating parameters related to the myogenic response of the afferent arteriole by arbitrary amounts to mimic scenarios of diminished autoregulation, which can impact blood delivery to the glomerular capillaries and, consequently, kidney filtration. Here, we aim to connect these parameters to the actual extent of GFR decline for clinical stages. A direct correlation between GFR and SNGFR is assumed under stable nephron population *n*:

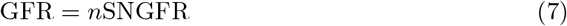

This relationship is fundamental for understanding renal function and dysfunction in the model’s context. However, pathological states may introduce complexities that alter this direct relationship.

To classify distinct stages of functional impairment based on the observed decline in GFR, as outlined in Table 2, we use the following ratios to calculate the SNGFR input to the model for diseased rat kidney stages *i* that correspond to the human disease stages:

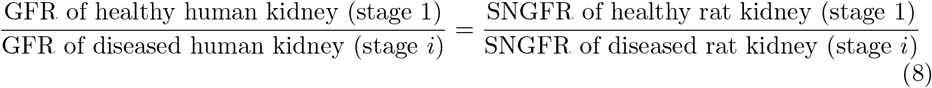

**Table 2.** Stages of CKD according to GFR ranges from [17, 18].

| Stage $i$ | GFR range (ml/min) | GFR mid-value used (ml/min) | SNGFR diseased rat kidney calculated (nl/min) |
| --- | --- | --- | --- |
| 1: healthy | $\text{GFR} \geq 90$ | 90 | 30 |
| 2: mild | $60 \leq \text{GFR} < 90$ | 74.5 | 24.833 |
| 3a: mild to moderate | $45 \leq \text{GFR} < 60$ | 52 | 17.333 |
| 3b: moderate to severe | $30 \leq \text{GFR} < 45$ | 37 | 12.333 |
| 4: severe | $15 \leq \text{GFR} < 30$ | 22 | 7.333 |
| 5: failure | $\text{GFR} \leq 15$ | 15 | 5 |

The derivation of Equation (8) assumes that the SNGFR for humans and rats is directly proportional. This proportionality remains consistent in analogous disease conditions. This relationship can be elaborated as follows. Let *hh* represent the healthy stage for humans and *hi* represent stage *i* for humans, where the number of human nephrons is constant (*n*_*h*_):

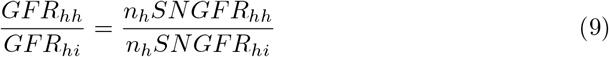

Similarly, let *rh* denote the healthy stage for rats and *ri* denote stage *i*, with the rat nephron number (*n*_*r*_) considered constant:

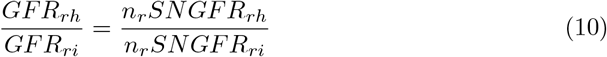

Assuming that the SNGFR for humans is a scalar multiple of the SNGFR for rats, then we can equate the left-hand side of Equation (9) for humans with the right-hand side of Equation (10) for rats to yield Equation (8). This allows for comparing GFR changes between healthy and diseased states across species.

The dynamics of fluid flow within the glomerulus are crucial to our understanding, as a portion of the afferent arteriole’s flow, *Q*_*A*_, contributes to the filtration process. We use the SNGFR data from different stages of chronic kidney disease to calculate *β* for each stage *i*:

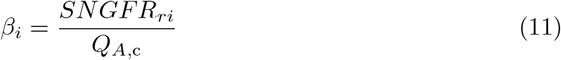

where *Q*_*A*,c_ is the volumetric flow rate (Equation (6)) evaluated at the control state *P* = 100 mm Hg (for the healthy case). We assume that *β* depends directly on the SNGFR, which is the variable changing in our model depending on disease stage *i*. The *β* parameter is pivotal as less of the blood is processed by glomerular filtration to remove wastes in more severe stages of CKD. This should reduce the flow rate *F* (*D*(*t*)) (Equation (5)) and thus advection in Equation (4). We have set the baseline *SNGFR*_*rh*_ at 30 L/min [10, 19]. *SNGFR*_*ri*_ values for all stages are listed in Table 2.

In [20], micropuncture experiments were conducted involving direct measurement of fluid in kidney tubules and linescan methods to measure the SNGFR. The findings from [20] indicate that the control rats have SNGFR values in the range 19.43–32.21 nl/min, which is consistent with our observed values for the healthy cases. For rats experiencing ischemia/reperfusion—indicative of acute kidney injury—the SNGFR values show a significant decrease to 5–19 nl/min on average. These experimental results align with the calculated SNGFR values (Table 2).

## 3 Results

A computational model simulating myogenic and tubuloglomerular feedback (TGF) mechanisms was used to investigate renal blood flow regulation across chronic kidney disease (CKD) stages the results obtained are described here. The model was coupled with varying glomerular filtration rates (GFR) at different CKD stages. The model’s output was analyzed to assess autoregulation under pressure inputs *P* between 60-180 mm Hg, compared to a 100 mm Hg control state. Pressure perturbations were step increases or decreases from initial pressures and steady-state solutions. In *in silico* trials for each disease stage evaluated the model’s autoregulation predictions and allowed comparisons of renal function under physiological and pathological conditions. Steady-state and dynamic responses to various pressures and their effects on afferent arteriole diameter, smooth muscle activation, single nephron GFR, and macula densa chloride ion Cl^−^ concentration were examined.

### 3.1 Afferent arteriole diameter and activation

In the simulations for afferent arteriole diameter, responses were evaluated under step-down pressures (100 to 60 mm Hg) and step-up pressures (100 to 180 mm Hg), providing insights into autoregulation across various stages of CKD.

For step-down pressures (Fig. 2), arteriole diameters *D*_*c*1_ increased progressively as pressure decreased, with healthy Stage 1 maintaining a control diameter of *D*_*c*1_ = 14.7 µm at 100 mm Hg. As pressures were reduced, the arteriole dilated, reflecting effective vasodilation and autoregulation to maintain stable blood flow. However, in later stages of CKD (Stages 3b–5), the ability to vasodilate was significantly impaired. This reduction in vasodilatory capacity suggests that, in these advanced disease stages, the autoregulatory mechanisms become insufficient to maintain proper renal blood flow as blood pressure decreases.

**Fig 2.**
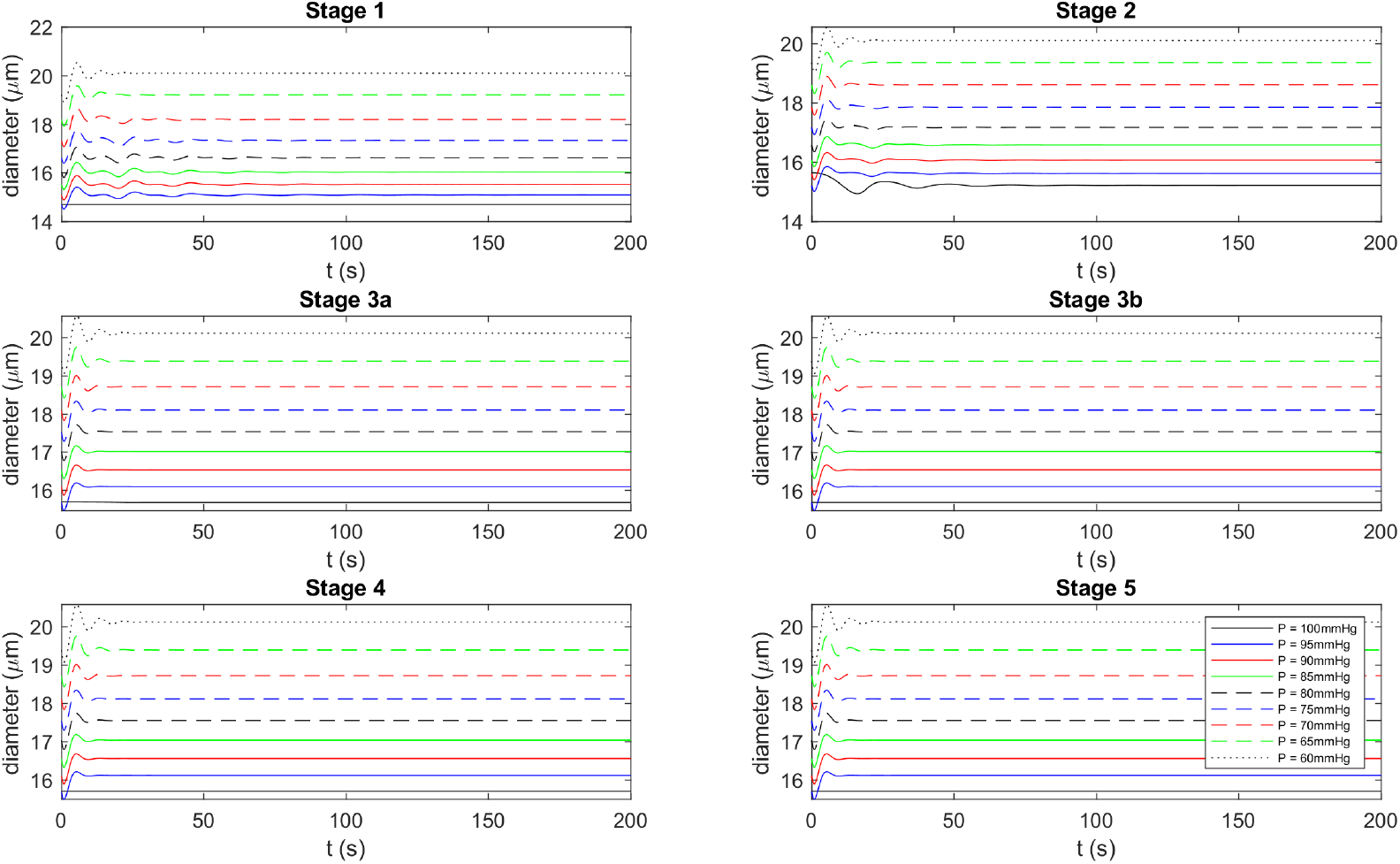
Arteriole diameter vs. time for step-down pressures (60–100 mm Hg) from the control state at 100 mm Hg. Note: the *y*-axis scales are not held constant across the stages.

Under step-up pressure conditions (Fig. 3), the afferent arteriole exhibited vasoconstriction in response to increasing pressures. In Stage 1, the arteriole responded with appropriate constriction, thereby protecting the kidney from excessive blood flow at higher pressures. However, in advanced CKD stages, the ability to constrict was significantly reduced, with the arteriole reaching its minimum diameter rapidly. This indicates a diminished capacity for autoregulation in response to elevated pressures, a critical problem in preventing damage to the kidney’s filtration units (nephrons) in advanced disease.

**Fig 3.**
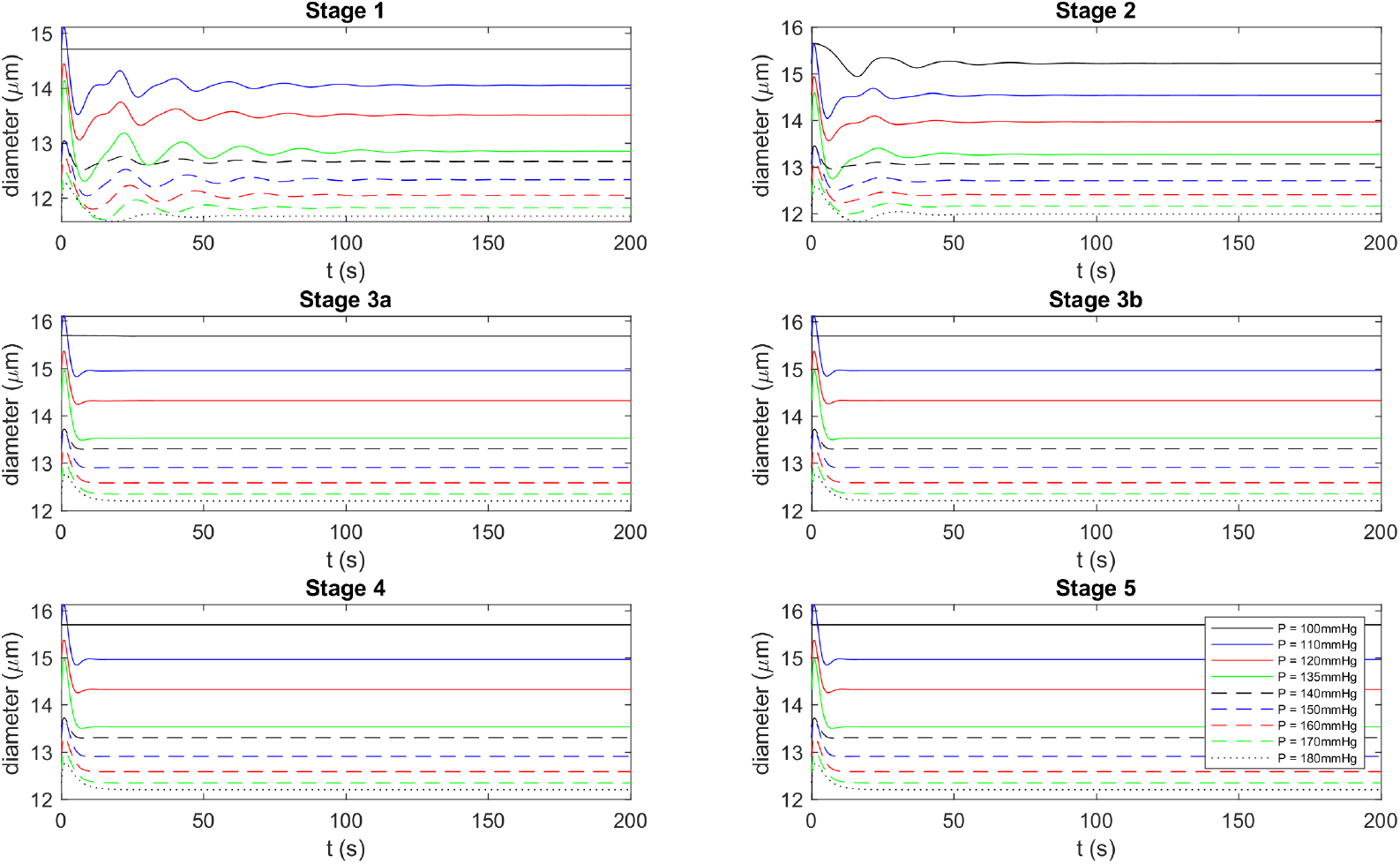
Arteriole diameter vs. time for step-up pressures (100–180 mm Hg) from the control state at 100 mm Hg. Note: the *y*-axis scales are not held constant across the stages.

In healthy and disease stages, the diameter of the afferent arteriole exhibits a decrease with an increase in perfusion pressure, a characteristic feature of autoregulation [21]. However, the arteriole diameters are consistently larger during pathological conditions compared to the healthy state at equivalent pressures. This discrepancy may be attributed to conditions such as hypertension or diabetes mellitus, which can impair the kidney’s autoregulatory mechanisms, thereby reducing the afferent arteriole’s ability to adjust its diameter in response to perfusion pressure changes. Chronic kidney disease can result in structural changes in the renal microcirculation, including conditions like glomerular sclerosis or arteriolar hyalinosis, which contribute to the reduction in the afferent arteriole diameter [22].

Steady-state responses across a range of pressures (60–180 mm Hg, Fig. 4) further highlight the impact of CKD progression on arteriole behavior. In the healthy state, the diameter remained consistent at 20.1 µm when the pressure was 60 mm Hg. However, as pressures increased, the diameter decreased, reflecting normal autoregulation. In contrast, for advanced CKD stages, the arterioles remained significantly more dilated at higher pressures, indicating an impaired ability to constrict. This loss of regulatory function is a hallmark of CKD progression, as the arterioles are no longer able to effectively manage renal blood flow at elevated pressures.

**Fig 4.**
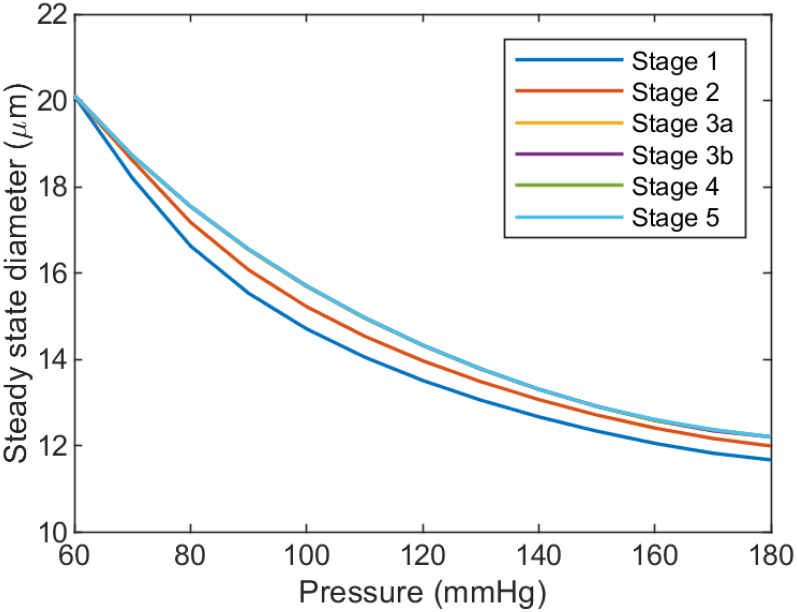
Steady-state arteriole diameter vs. pressure ranges (60–180 mm Hg). Note that the values for Stages 3a–5 are indistinguishable from one stage to another. factor in the progression of CKD.

In a normal kidney, the vascular smooth muscle activation constricts in response to increased pressure in the vascular wall, resulting in a gradual decrease in its diameter across a pressure range of 60–180 mm Hg [23]. This decrease in vessel diameter and increase in activation (Fig. 5) is an expected behavior of the myogenic response and is the primary regulator that leads to autoregulation of blood flow in the kidney [8]. However, in kidneys affected by diseases such as diabetes, the ability of the afferent arteriole to respond to pressure changes is compromised, leading to a significant reduction in its ability to constrict. This impaired response to pressure changes is a key

**Fig 5.**
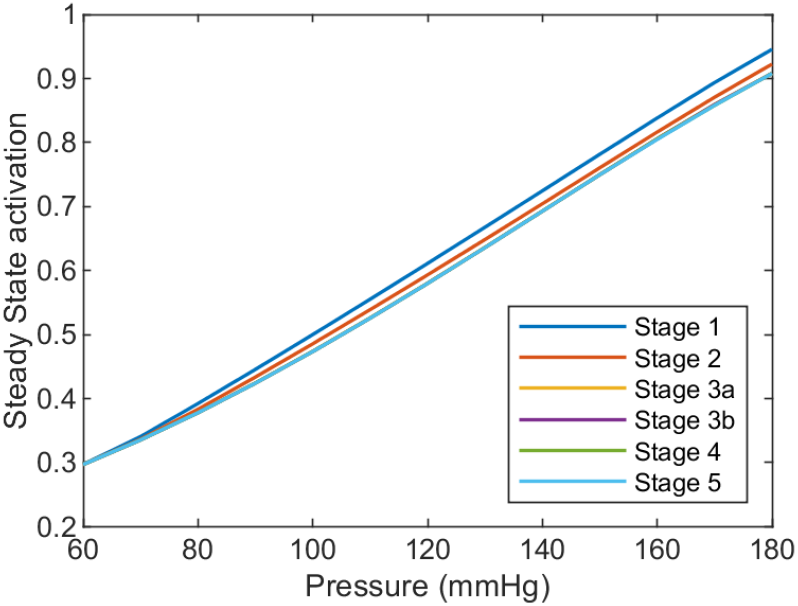
Steady-state arteriole activation vs. pressure ranges (60–180 mm Hg). Note that the values for Stages 3a–5 are indistinguishable from one stage to another.

The model predictions were validated against experimental data from studies on afferent arteriole behavior under varying pressure conditions. The findings from Sanchez et al. (1989) [24], which investigated the effect of perfusion pressures (80–180 mm Hg) on rat afferent arterioles, demonstrated vasoconstriction by 16.4% when pressure was raised from 80 to 120 mm Hg. These experimental results align well with our model, confirming the predicted myogenic response and supporting the model’s representation of autoregulatory mechanisms.

Similarly, Loutzenhiser et al. (2002) [25] showed a graded reduction in arteriole diameter as renal arterial pressure increased, particularly beyond 80 mm Hg. Our model’s predictions closely followed the experimental findings, demonstrating both narrowing of the arteriole at high pressures and a comparable quadratic relationship between pressure and arteriole diameter. The consistency of the model’s output with experimental data across various pressure conditions validates its ability to simulate renal autoregulation effectively.

### 3.2 Single nephron glomerular filtration rate

In simulations examining the impact of CKD progression on single nephron glomerular filtration rate *SNGFR* = *Q* = *βQ*_*A*_, step-down and step-up pressure perturbations were applied to assess how well SNGFR is maintained across different CKD stages. In both cases, the objective is to reach the desired *SNGFR*_*i*_ value for each stage, independent of the pressure value. As disease stages progress, the variation required to attain the initial state value decreases across different pressure states. This phenomenon may be attributed to the functioning of the kidney.

For step-down pressures (60–100 mm Hg), as shown in Fig. 6, SNGFR progressively decreases in all stages. In healthy Stage 1, the kidney quickly stabilizes the SNGFR, indicating functional autoregulation. However, in more advanced CKD stages (Stages 3b–5), the kidney’s ability to stabilize the SNGFR diminishes. This indicates a loss of autoregulatory efficiency, with filtration rate dropping significantly in response to reduced pressures.

**Fig 6.**
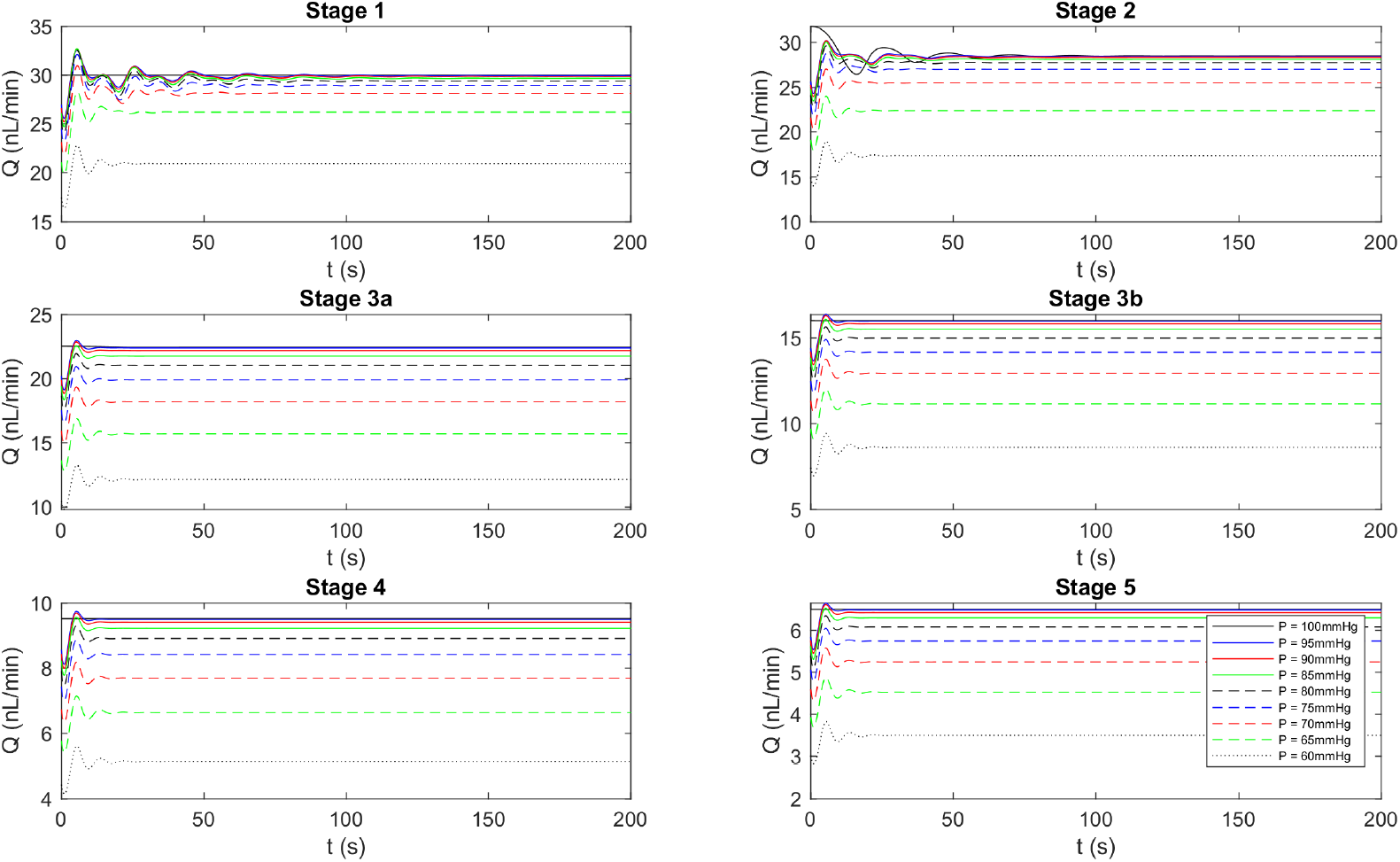
Flow rate (*Q*) vs. time for step-down pressures (60–100 mm Hg) from the control state 100 mm Hg. Note: the *y*-axis scales are not held constant across the stages.

For step-up pressures (100–180 mm Hg), illustrated in Fig. 7, the SNGFR increases in all stages as pressure rises. In healthy kidneys (Stage 1), the SNGFR initially rises with pressure but stabilizes at a certain threshold, indicating effective autoregulation. However, in advanced CKD stages (3b–5), the SNGFR continues to rise without reaching a steady state. This failure to stabilize suggests impaired autoregulation in later stages of CKD, leaving the kidney more vulnerable to damage from sustained elevated pressures.

**Fig 7.**
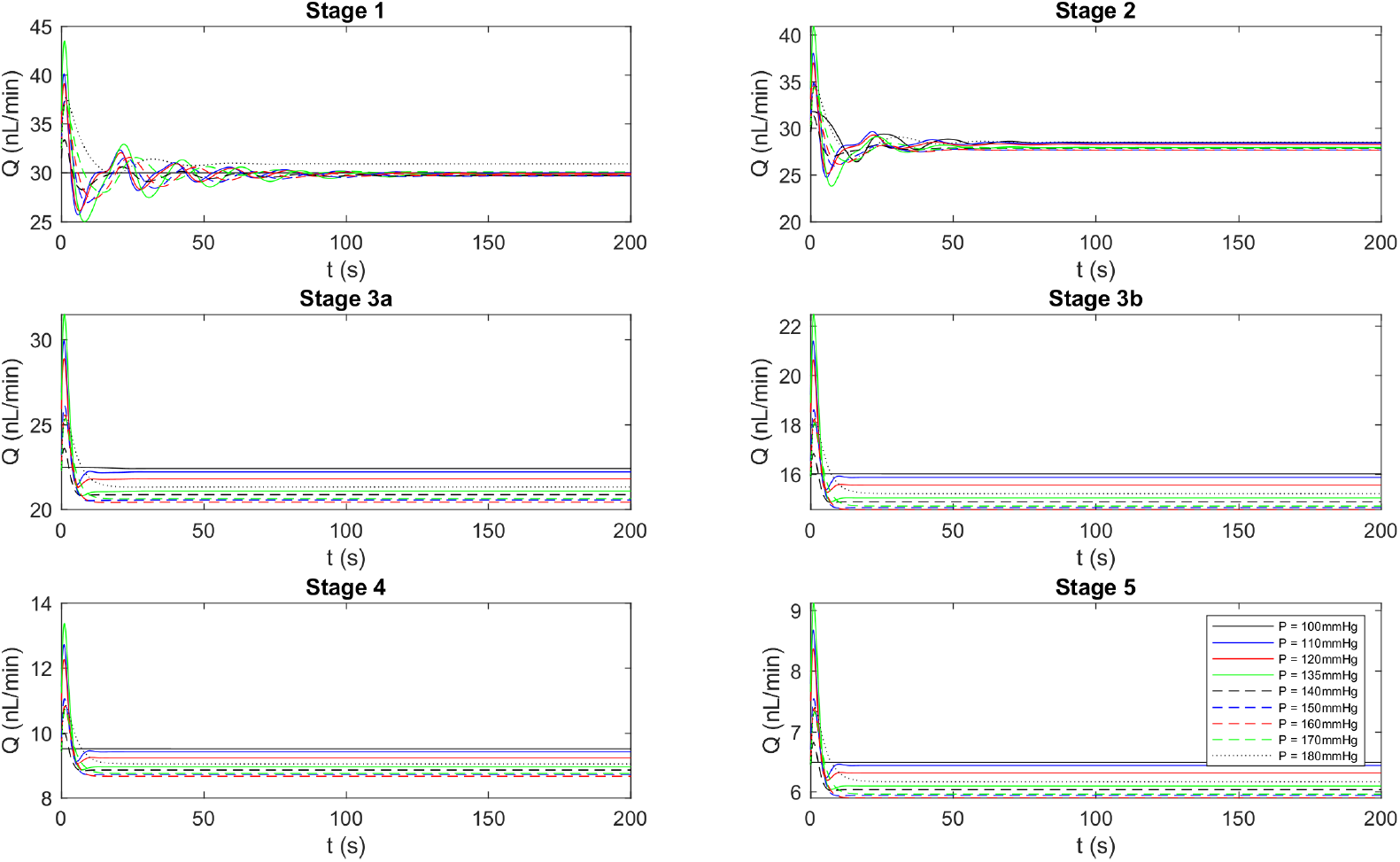
Flow rate vs. time for step-up pressures (100–180 mm Hg) from the control state 100 mm Hg. Note: the *y*-axis scales are not held constant across the stages.

#### 3.2.1 Model validation

In Fig. 8, the relationship between arteriolar pressure and the fluctuation in normalized SNGFR in kidneys highlights how a normal kidney’s autoregulatory mechanism maintains stable SNGFR despite pressure changes in Stage 1. Note that normalized SNGFR is *Q*_*A*_*/Q*_*A*,c_, which is independent of *β*_*i*_. Fig. 8 also illustrates the implications of impaired autoregulation, which can lead to kidney damage. In healthy kidneys, an increase in arteriolar pressure leads to a corresponding rise in SNGFR up to a certain point (80 mm Hg). Beyond this, the SNGFR stabilizes in Stage 1, showcasing the kidney’s capacity to counteract pressure changes and maintain a consistent filtration rate. As arteriolar pressures exceed the 100 mm Hg threshold, Stages 3a–5 deviate from the anticipated pattern that is to maintain a constant filtration. This deviation suggests a potential impairment in the autoregulatory mechanism, as evidenced by the fluctuations in normalized SNGFR. When one kidney is slowly deteriorating, the remaining kidney increases its SNGFR to compensate. Despite pressure variations, the SNGFR stays relatively stable, indicating the kidney can adjust its resistance and protect the glomeruli from high pressure (Stage 2).

**Fig 8.**
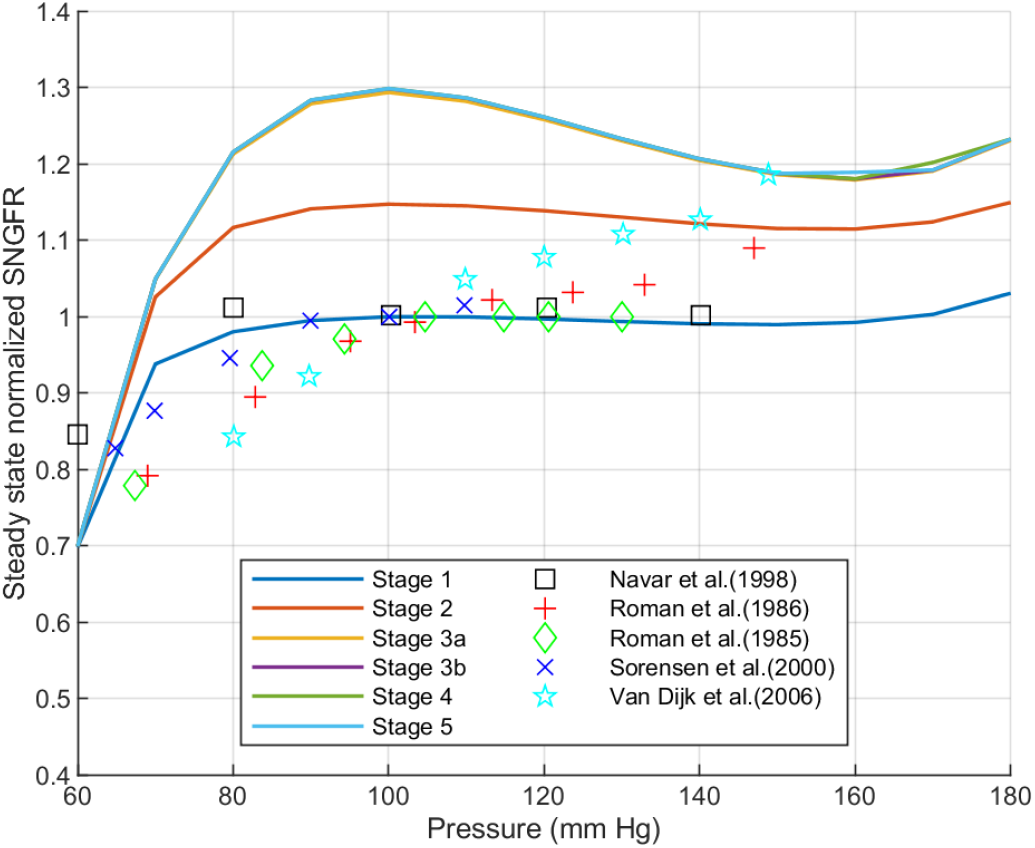
Steady-state SNGFR vs. pressure. Model-predicted values of various stages of normalized flow in the afferent arteriole are plotted against pressures in the 60–180 mm Hg range to observe the relationship between steady-state SNGFR and pressure. The model outcomes for the healthy stage compared with empirical findings from rat experiments: [26–30].

In Stages 3a–5, it is observed that the SNGFR continues to rise with blood pressure up to 100mm Hg; without reaching a steady state in these stages, there are fluctuations. This suggests that the kidney’s capacity to manage its resistance is compromised, making it vulnerable to hypertension. This could potentially lead to injuries in the glomerulus and subsequent fibrosis. In the final stages, this can result in damage to the glomerulus. The outcomes depicted in Fig. 8 are supported by the model presented in the work of [31]. Their study compared the stages of the disease (glomerulosclerosis, diabetes, hypertension) with healthy individuals. The similarities between their findings and our results in Fig. 8 provide validation for our model.

The description provided here is consistent with the expected behavior of a healthy kidney compared to a diseased one. A healthy kidney typically can autoregulate and maintain a stable SNGFR despite changes in pressure. On the other hand, kidneys affected by disease may lose this autoregulation capability, leading to variations in the SNGFR.

For the healthy stage, the model-predicted values of normalized SNGFR are represented in Fig. 8. This figure showcases the effects of variation of arterial pressure in the 60–180 mm Hg range. To provide a comprehensive analysis, the model results are compared with experimental data values obtained from autoregulation studies conducted in rats [26–30]. Note that normalization is relative to stage 1 at 100 mm Hg.

In [31], an illustration is presented that demonstrates the autoregulation of renal blood flow in healthy and hypertensive individuals. The model in [31] compares the relationship between systolic blood pressure and normalized SNGFR under different conditions, including but not limited to normal, diabetes, fawn hooded hypertensive (FHH) rats, spontaneously hypertensive rats (SHR), and low/high salt intake. When comparing the plots presented from the publications [31, 32] with our model, particularly across various stages of the disease, our model consistently agrees with the experimentally validated findings reported in the published research [31, 32]. This suggests that autoregulation is effectively maintained during the early stages of the disease. However, for the subsequent stages, starting from Stage 3a, there appears to be a noticeable impairment in autoregulation.

### 3.3 Chloride ion concentration at the macula densa

The TGF mechanism, the main role in regulating GFR, heavily relies on the Cl^−^ concentration at the macula densa (CMD). The macula densa consists of cells that function as salt sensors, detecting the Cl^−^ concentration in the filtrate. A decrease in blood volume can lead to a drop in blood pressure, resulting in reduced CMD. These changes are detected by the macula densa cells [33]. The lower blood volume, accompanied by decreased pressure, enters the afferent arteriole and, subsequently, the glomerulus, which results in decreased GFR—this filtration rate is the main factor that the macula cells are designed to detect. An important thing to be noted from this information that is related to the model is that as the GFR reduces, less fluid is filtered from the blood into the nephron, leading to a decrease in the Cl^−^ concentration in the tubular fluid [34].

As can be seen in Fig. 9, in this model, Stage 1 (representing normal kidney) exhibits the highest CMD, suggesting a robust Cl^−^ concentration gradient at the macula densa, consistent with normal kidney function.

**Fig 9.**
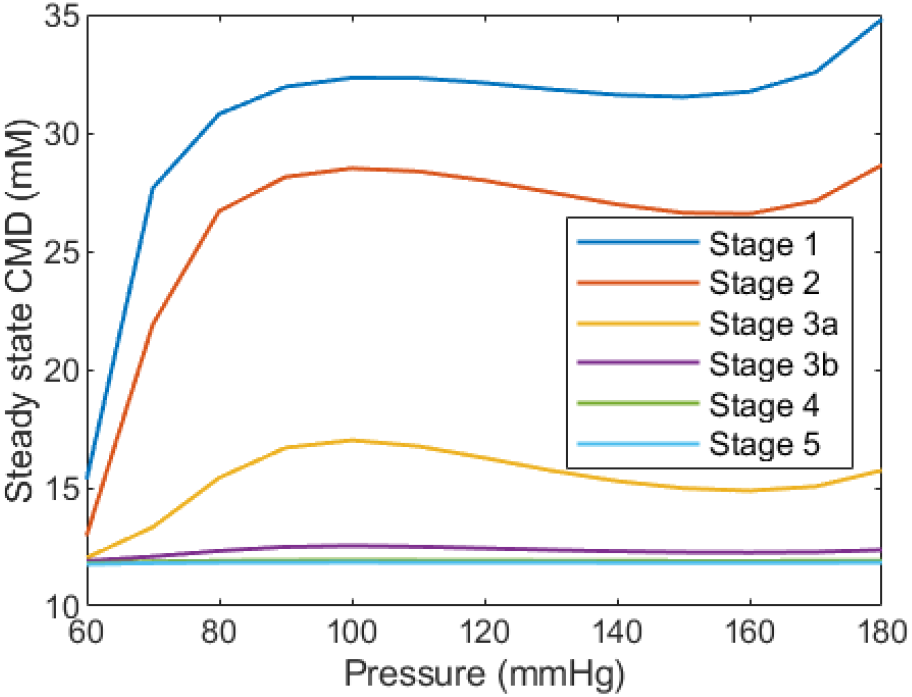
Steady-state chloride ion Cl^−^ concentration at the macula densa (CMD) vs. pressure.

As the stages progress from Stage 2 to Stage 5, there is a noticeable decrease in CMD, indicating a decline in kidney function. The highest stages (e.g., Stage 5) represent more advanced conditions of kidney impairment, with a considerable drop in CMD, which may be indicative of an advanced disease state.

During Stage 2, there is an initial incline in CMD followed by a consistency in the curve with minimal fluctuation, reflecting a change in SNGFR. This could be interpreted as the kidney’s attempt to adapt and partially restore the CMD towards healthier levels, although it does not fully achieve the CMD observed in Stage 1. This pattern may represent the kidney’s intrinsic ability to counteract fluctuations in SNGFR to some extent. In contrast, Stage 5, representing an advanced disease state, shows that CMD levels off quickly after the initial change and does not exhibit the fluctuations seen in the earlier stages (Fig. 9). This could suggest that in more severe stages of kidney disease, there is a loss of the dynamic regulatory capacity, and the system settles into a new, lower CMD without attempting to modulate it further, reflecting a diminished or exhausted physiological response. A consistently low level of Cl^−^ might indicate a problem with sodium reabsorption in the kidney, potentially leading to conditions like hyponatremia (low blood sodium levels).

Figs. 10–11 illustrate the changes in CMD over time response to step-down and step-up changes in pressure from a control state at 100 mm Hg. CMD reaches a steady-state value after an initial transient response to the step-down in pressure (Fig. 10). This steady-state value varies across the different stages, with Stage 1 having the highest CMD and Stage 5 having the lowest CMD. The time required to reach the steady-state CMD appears to be longer for the earlier stages (Stages 1–2) compared to the later stages (Stage 3b–5). This could be an indication of the kidney’s adaptive mechanisms attempting to restore the CMD to higher levels in the earlier stages, as described previously for Fig. 9. Overall, even in step-up pressures (Fig. 11), the earlier stages (Stages 1–3a) have larger fluctuation before reaching steady states. This gives the idea that the kidney’s autoregulatory capacity is more diminished in the later stages.

**Fig 10.**
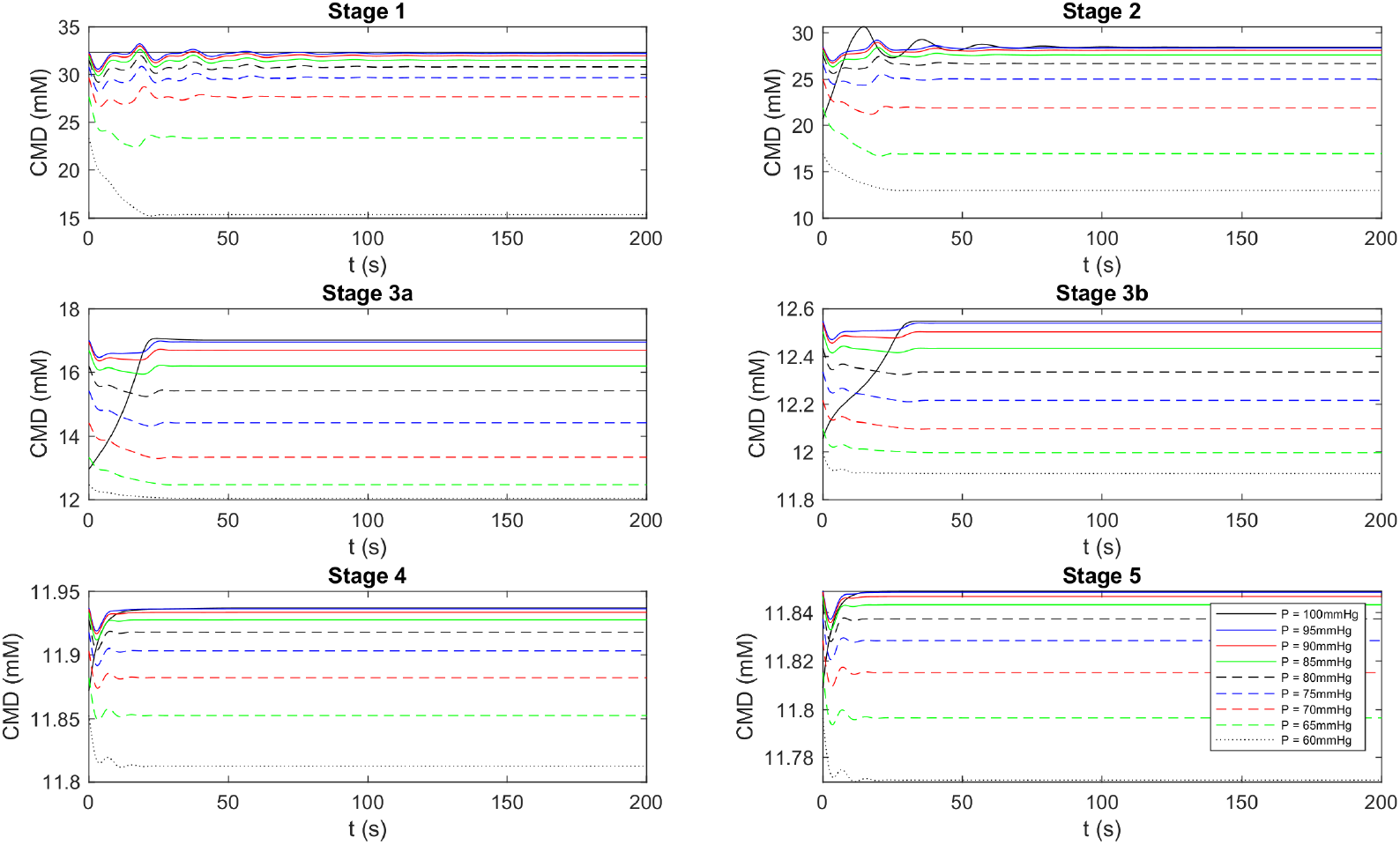
Chloride ion Cl^−^ concentration (CMD) at the macula densa vs. time for step-down pressures (60–100 mm Hg) from control state 100 mm Hg. Note: the *y*-axis scales are not held constant across the stages.

**Fig 11.**
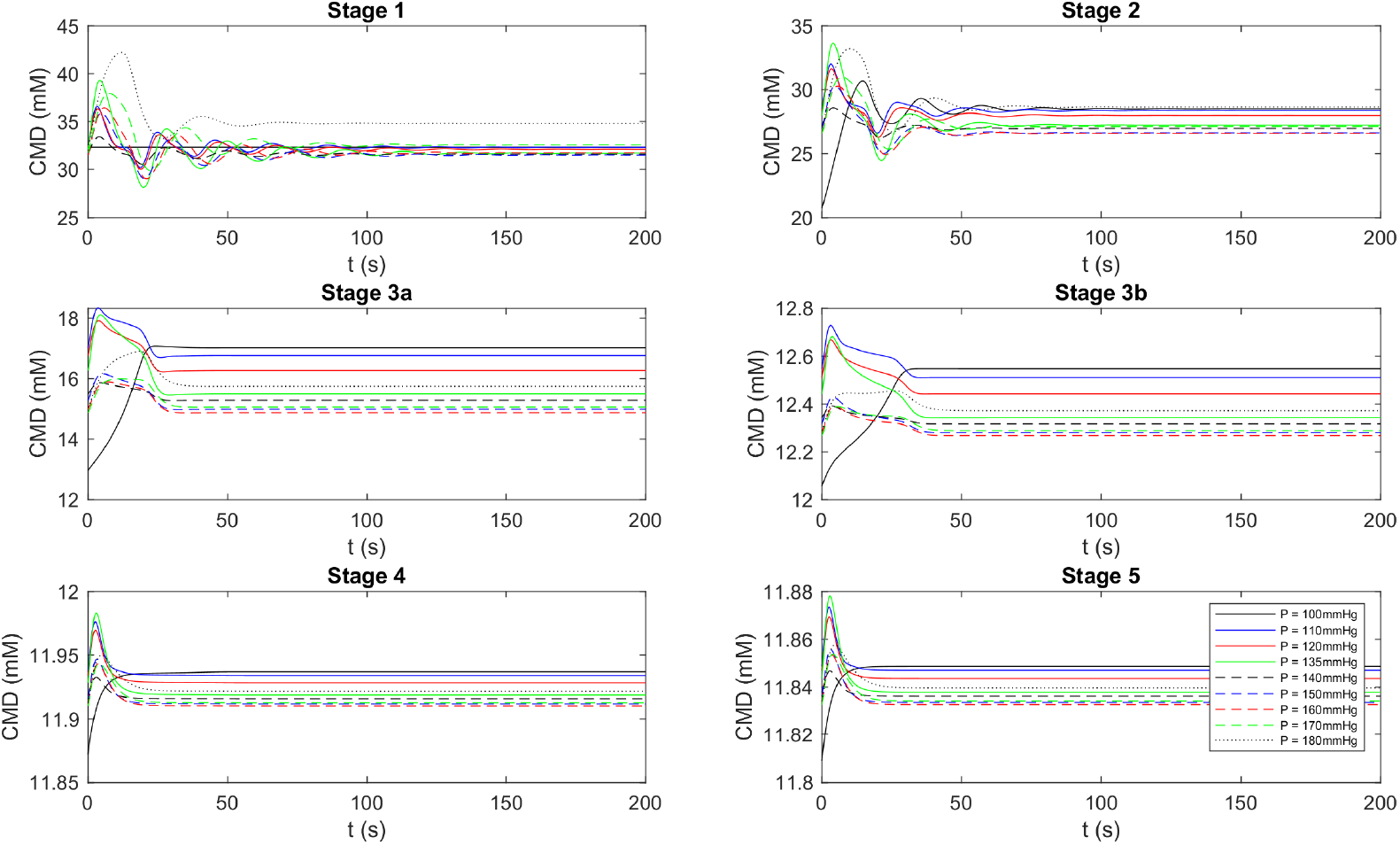
Chloride ion Cl^−^ concentration at the macula densa (CMD) vs. time for step-up pressures (100–180 mm Hg) from control state 100 mm Hg. Note: the *y*-axis scales are not held constant across the stages.

The study by [32] elucidates the mechanisms by which the body maintains electrolyte and fluid balance in both healthy and diseased states. Under normal physiological conditions, Cl^−^ is reabsorbed in the TAL through the Na-K-2Cl cotransporter. This process is facilitated by the basolateral Na-K-ATPase, which maintains a low intracellular sodium concentration. Consequently, a high osmotic gradient is established in the medulla, allowing for urine concentration. Burnier [32], demonstrated that sodium excretion through urine remains unaffected by changes in blood pressure in individuals with normal blood pressure, suggesting a blood pressure that is resistant to salt and can maintain sodium balance without altering blood pressure.

Conversely, hypertensive patients with normal kidney function exhibit a higher rate of sodium excretion through urine compared to those with normal blood pressure at the same blood pressure level, implying that these hypertensive patients have blood pressure that is sensitive to salt, requiring a higher pressure for the excretion of the same amount of sodium. Moreover, hypertensive patients with impaired kidney function demonstrate a lower rate of sodium excretion through urine compared to individuals with normal blood pressure at the same blood pressure level. This finding suggests that these patients have both salt-sensitive blood pressure and impaired renal function, leading to a reduced capacity for sodium excretion and an increased susceptibility to sodium and fluid overload.

At the terminal end of the TAL, where the macula densa is located, salt concentrations depend primarily on the flow rate. In the physiological flow range of *≈* 5–30 nl/min of SNGFR [35], it is estimated that Na^+^ and Cl^−^ concentrations vary by *≈* 30–40 mM, with low values of *≈* 10–15 mM, which aligns with the Cl^−^ concentrations observed in Fig. 9 (highest *≈* 30 mM and lowest *≈* 13 mM). In Stages 3–5 of CKD, alterations in Cl^−^ transport within the TAL can arise due to tubular dysfunction, changes in TGF, and the effects of therapeutic interventions. However, the specific modifications in Cl^−^ handling within the TAL are not explicitly delineated in [32]. With kidney dysfunction and the loss of functioning nephrons, there is an increased potential for Cl^−^ retention, which can contribute to volume overload and hypertension.

## 4 Discussion

In general, for the plots between diameter, Cl^−^ concentration, and SNGFR vs. time an elevation in pressure is linked to a relatively minor rise in glomerular filtration, which in turn, amplifies the delivery of filtrate to the nephron, resulting in enhanced urine production [36]. A pivotal factor behind the occurrence of oscillatory responses in the kidney model is the autoregulatory mechanism. This physiological ability enables the body to uphold consistent blood flow to organs despite fluctuations in arterial pressure. This process involves the modulation of arteriolar diameters through vasodilation and vasoconstriction, which aids in fine-tuning resistance and maintaining blood flow within a specific range. Oscillatory responses arise as arterioles continuously adapt their diameters in an attempt to stabilize blood flow against pressure fluctuations. Additional contributors to these oscillations include the myogenic and TGF mechanisms. The regulation of blood flow and pressure involves intricate interactions between multiple feedback loops, yielding oscillatory behavior in arteriolar diameter.

In all the scenarios outlined, the plots capture the temporal changes in factors such as flow concentration. Notably, for the diseased state, a consistent alteration pattern persists across the entire process, irrespective of the pressures applied. This observation underscores the abnormal nature of the diseased kidney, as it lacks the oscillatory responses characteristic of a healthy kidney, considering the understood mechanisms. Similarly, in cases where oscillations are externally induced, such as modifying the SNGFR to 30 [10], 24.833, 17.333, 12.333, 7.333, and 5 nl/min (Table 2), the resulting plots reveal the absence of fluctuations or oscillatory responses.

These results can be helpful in further analysis. Arciero et al. [8] discussed how the observed constriction in Fig. 4, i.e., decrease in vessel diameter with the expected behavior of the myogenic response and is the primary regulator that leads to autoregulation of kidney blood flow. Epp et al. [37] described how changes in diameter at the capillary level enable a very localized regulation of blood flow but does not provide any information on how these changes occur over time.

## 5 Conclusions

The primary objective of this computational endeavor was to elucidate the mechanisms by which the kidney maintains homeostasis of blood flow and glomerular filtration rate despite variations in arterial pressure in disease conditions. The functioning of kidneys, particularly under diseased conditions such as CKD, which includes conditions like diabetic nephropathy and glomerular damage, on renal blood flow regulation was investigated. This research focused on understanding how variations based on the glomerular filtration rate impact key kidney function outputs, such as afferent arteriolar diameter, smooth muscle activation, and the chloride ion Cl^−^ concentration at the macula densa. We have analyzed these factors by considering the dynamics of the mathematical model of ODEs/PDE to study blood flow control in the kidney, which has provided new insights into the maintenance of autoregulation.

The analysis in the results revealed that as CKD progresses from early to advanced stages, afferent arteriole diameter demonstrated a decrease in diameter with increasing renal arterial pressure. Notably, even in pathological conditions, the diameter decreased but remained slightly elevated compared to a healthy kidney state. Plots depicting the changes in diameter over time indicated a stabilization of the diameter after a certain period, with a quicker stabilization observed in the advanced disease stages compared to the healthy stage. Furthermore, an increase in smooth muscle activation was observed with rising pressure. In the initial stages, autoregulation exhibited dynamic behavior, reaching a stable constant value after a certain period. However, no fluctuations were observed in the final stages, deviating from the normal behavior seen in stage 1, suggesting impaired autoregulation in the later stages of kidney disease.

For different stages of GFR, Fig. 8 demonstrates how SNGFR changes in response to variations in pressure levels. Stage 1 maintains autoregulation, as it is constant throughout the range of pressure variations, and the GFR decreases the blood flow rate and exhibits distinct patterns of pressure. The data points with markers include experimental data points from rat studies; the comparison can validate the accuracy of the mathematical model in simulating SNGFR responses to pressure perturbations. The chloride concentration at the macula densa, which plays a crucial role in the tubuloglomerular feedback mechanism, also decreases progressively with advancing CKD stages, reflecting the diminished ability of the kidney to regulate sodium and fluid balance. Overall, the study provides valuable insights into the hemodynamic dysregulation accompanying CKD progression, paving the way for a better understanding of the underlying mechanisms.

The current research has provided valuable insights into the hemodynamic changes that occur in CKD. However, further study requires a comprehensive understanding of the complex factors involved in renal autoregulation and its impairment in disease states. Future investigations could expand the mathematical model to incorporate additional factors such as hormonal regulation, including angiotensin II and endothelin, and their effects on renal hemodynamics. Using the virtual kidney model, the impact of renin-angiotensin system (RAS) and angiotensin-converting enzyme (ACE) inhibitors on the modeled parameters could be studied. This would allow for an evaluation of their effectiveness in restoring autoregulatory mechanisms. The model could also be extended to investigate the long-term consequences of impaired autoregulation on the decline of renal function and the progression of CKD. By addressing these areas, future research could enhance our understanding of the complex pathophysiology of CKD and potentially contribute to the development of more effective therapeutic strategies for managing this debilitating condition

## Acknowledgments

This work was supported by the National Science Foundation CAREER grant 2133411 and resources from the University at Buffalo. The content is solely the responsibility of the authors and does not necessarily represent the official views of the funding agency. We also acknowledge Dr. Rudiyanto Gunawan, who served as a committee member for B. Maddineni’s M.S. thesis.

